# A CRISPR/Cas9-based system using dual-sgRNAs for efficient gene deletion in *Mycobacterium abscessus*

**DOI:** 10.1101/2025.05.28.656731

**Authors:** Linai Li, Dan Wang, Xin Li, Yuxiang Hu, Shengjuan Bao, Taibing Deng, Qinglan Wang

## Abstract

The increasing global prevalence of *Mycobacterium abscessus* infections presents a significant clinical challenge due to the pathogen’s intrinsic resistance to multiple antibiotics and poor treatment outcomes. Despite the necessity of genetic tools for studying its physiology, pathogenesis, and drug resistance, efficient methods for large-fragment deletions remain underdeveloped. Here, we report a CRISPR/Cas9-based dual-sgRNA system employing *Streptococcus thermophilus* CRISPR1-Cas9 (Sth1Cas9), enabling efficient large-fragment knockout in *M. abscessus* with deletion efficiencies exceeding 90% at certain loci and spanning up to 16.7 kb. Furthermore, we systematically optimized the modular arrangement of genetic components in Cas9/dual-sgRNA expression plasmids and refined their construction workflow, achieving a significant reduction in cassette loss rates while enabling single-step plasmid assembly. Notably, deletion efficiency was position-dependent rather than correlated with target size, suggesting an influence of chromatin structure on editing outcomes. As the first CRISPR/Cas9-based platform capable of kilobase-scale deletions in *M. abscessus*, this system advances functional genomics studies and facilitates targeted investigations into virulence and antibiotic resistance mechanisms.

## 1. Introduction

*Mycobacterium abscessus* has emerged as a critical global public health threat, particularly among immunocompromised individuals (1). As a rapidly growing nontuberculous mycobacterium (NTM), it causes aggressive pulmonary, cutaneous, and disseminated infections, with clinical treatment failure rates exceeding 50% (2). This high failure rate is largely attributed to its multidrug-resistant (MDR) phenotype, including intrinsic resistance to first-line tuberculosis drugs (e.g., isoniazid, rifampin) and inducible macrolide resistance mediated by the *erm(41)* gene (3).

Furthermore, *M. abscessus* employs biofilm formation and morphotype switching (smooth [S] to rough [R]) to evade host immune clearance, leading to chronic relapsing infections. Surveillance data from the World Health Organization indicate a 3%–5% annual increase in global infection rates, with colonization rates approaching 15% in cystic fibrosis patients, underscoring the urgent need for improved genetic tools to facilitate targeted therapeutic strategies.

Deciphering the biology, pathogenesis, and drug resistance mechanisms of *M. abscessus* necessitates efficient genetic tools. Traditional gene-editing approaches, such as homologous recombination, exhibit low efficiency (10^-6 to 10^-4) and are further hampered by high background antibiotic resistance, making mutant screening arduous (4-6). The construction of a single-gene knockout strain typically requires screening hundreds to thousands of colonies (6), severely limiting functional genomics research. While some studies attempted to mitigate background noise using fluorescent markers (e.g., tdTomato) (7), these modifications failed to fundamentally improve efficiency. Although transposon-based mutagenesis enables genome-wide screening, it lacks precision (6).

Recent advances in CRISPR/Cas9 technology offer a transformative alternative for genetic manipulation in *M. abscessus*. CRISPR/Cas9 employs a single guide RNA (sgRNA) to direct Cas9-mediated DNA double-strand breaks (DSBs), inducing frameshift mutations or small indels via non-homologous end joining (NHEJ) (8-12). The *Streptococcus thermophilus* CRISPR1-derived Cas9 (Sth1Cas9) has demonstrated high editing efficiency in *M. abscessus*, achieving a 10^2–10^4-fold improvement over conventional methods (8, 10). Prior studies implemented a dual-plasmid system to mitigate Cas9 cytotoxicity and incorporated fluorescent markers (e.g., mCherry) to enhance screening accuracy (10). However, while frameshift mutations effectively inactivate most genes, they may fail to disrupt genes with multiple translation initiation sites and can yield aberrant truncated proteins with unknown physiological consequences. Complete gene deletion is therefore necessary for certain applications.

Here, we report the development of an Sth1Cas9-dual-sgRNA system for highly efficient large-fragment gene knockout in *M. abscessus*. Our approach achieves deletion efficiencies exceeding 90% for some targets, with maximal deletion lengths of 16.7 kb. Notably, knockout efficiency does not correlate with fragment size, suggesting potential genomic location-dependent effects.

## 2. Materials and Methods

### 2.1 Bacterial Strains and Growth Conditions

*Mycobacterium abscessus* ATCC 19977 was used as the parental strain for all genetic manipulations. Cultures were grown in Middlebrook 7H9 broth or on Middlebrook 7H10 agar (BD Biosciences, USA) supplemented with 10% OADC (oleic acid, albumin, dextrose, catalase), 0.2% glycerol (Sangon Biotech), and 0.05% Tween-80 (Sangon Biotech, broth only) at 37°C. Plasmid-containing strains were maintained with the following antibiotics: kanamycin (MCE, USA; 100 μg/ml) for pCas9-mScarlet-containing strains and Zeocin (Invitrogen™, 20 μg/ml) for pQL033-X-sg-harboring strains. Anhydrotetracycline (aTc; MCE, 500 ng/ml) was used to induce CRISPR components. *Escherichia coli* DH5α (Tsingke; transformation efficiency >10^9^ cfu/μg) was used for plasmid construction and propagation in LB medium (Sangon Biotech) with kanamycin (100 μg/ml) or Zeocin (20 μg/ml) as required.

### 2.2 Plasmid Construction

To minimize toxicity, the CRISPR/Cas9 system was implemented using two separate plasmids.

The Cas9 expression plasmid was derived from pLJR962 (Addgene #115162) via site-directed mutagenesis to introduce A9D and A599H mutations, restoring nuclease activity and generating pCas9. The mScarlet fluorescent reporter was inserted into the EcoRV(New England Biolabs, USA; 10000 U/ml) site of pCas9 generate pCas9-mScarlet plasmid, allowing visual selection of transformants.

For targeted gene deletion, we developed a dual-sgRNA system based on pKM461 (Addgene #108320). As shown in Fig.S1, the anhydrotetracycline-inducible promoter (P_tet_), kanamycin resistance gene, and Sth1 sgRNA scaffold (Sth1SC) were amplified from pLJR962 and assembled into pKM461through SapI(New England Biolabs, USA; 10000 U/ml) and EcoRv sites, generating the pQL033-SapItwin-sgSC plasmid, which incorporated SapI restriction sites for Golden Gate assembly of target-specific sgRNAs. The pUC19-CS-ZeoR plasmid was constructed by inserting the Sth1SC-ZeoR-Ptet cassette into the pUC19 backbone. Target-specific protospacer sequences were designed to flank the genomic region of interest (e.g., Mab_0673/0674) and were incorporated into primers used for amplifying the Sth1SC-ZeoR-Ptet cassette, which contained SapI recognition sites. The resulting PCR product was then cloned into pQL033-SapItwin-sgSC to generate the final dual-sgRNA expression plasmid. All plasmids were confirmed by Sanger sequencing(Tsingke) before use.

### 2.3 Generation of Knockout Mutants in *M. abscessus*

Genetic manipulation of *M. abscessus* was performed using optimized electroporation protocols.

Competent cells were prepared by growing cultures to mid-log phase (OD600 ∼0.8-1.0) in 7H9 medium containing 0.05% Tween-80, followed by extensive washing with ice-cold 10% glycerol. For initial transformation, 1-2 μg of pCas9-mScarlet plasmid DNA was electroporated using a Bio-Rad Gene Pulser Xcell with parameters set to 2.5 kV, 25 μF, and 1000 Ω. Transformants were recovered in 7H9/OADC medium for 4 hours before plating on selective 7H11/OADC agar containing 100 μg/ml kanamycin. After 4-6 days incubation at 37°C, pink colonies expressing mScarlet were picked for verification by PCR and sequencing of the Cas9 cassette.

To generate knockout mutants, competent cells of the pCas9-mScarlet strain were transformed with 1 μg of the appropriate pQL033-X-sg plasmid (expressing dual sgRNAs). Following electroporation and recovery, transformants were selected on 7H11 plates containing both kanamycin (100 μg/ml) and Zeocin (20 μg/ml). For gene deletion, positive clones were grown to OD600 ∼0.8 in 7H9 medium with antibiotics, then split into two cultures - one induced with 500 ng/ml aTc and one uninduced control. After overnight induction, serial dilutions were plated on selective media with or without aTc to quantify survival rates. Potential knockout mutants were screened by PCR using primers flanking the target region, with successful deletions identified by the appearance of a smaller amplicon compared to wild-type. The deletion boundaries were confirmed by Sanger sequencing of PCR products.

## 3. Results

### 3.1 Design of a Dual-sgRNA CRISPR/Cas9 System for Gene Deletion in *M*.*abscessus*

To establish a CRISPR-based gene knockout system in *M. abscessus*, we engineered a dual-plasmid system comprising a Cas9 expression vector (pCas9-mScarlet) and a dual-sgRNA expressing plasmid (pKMZeoR-sg). The pCas9-mScarlet construct integrates into the L5 attB locus of the *M. abscessus* genome and carries an anhydrotetracycline (aTc)-inducible Sth1Cas9 gene, a kanamycin resistance marker, and an mScarletfluorescent reporter for visual selection of transformants (Fig. 1B). Following electroporation into wild-type *M. abscessus*, approximately 30% of kanamycin-resistant colonies lacked fluorescence, indicating a high false-positive rate on kanamycin plates.

**Figure 1.**
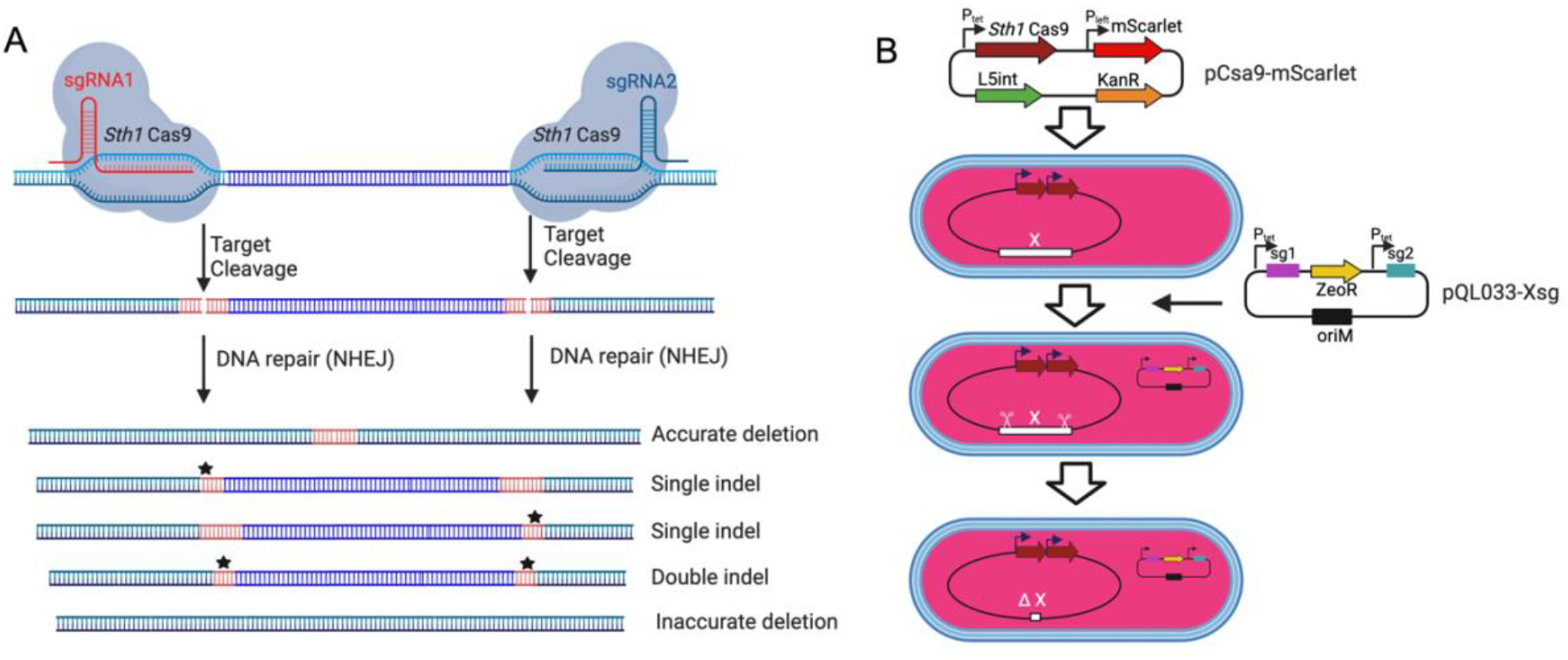
CRISPR-Cas9-mediated gene deletions using dual sgRNAs in *M. abscessus*. (A) Schematic of CRISPR-Cas9-induced gene deletions using dual sgRNAs. *M. abscessus* cells harboring pCas9-mScarlet and pQL033-Xsg plasmids express Sth1Cas9 and dual sgRNAs upon aTc induction. The sgRNAs guide Sth1Cas9 to the target genomic loci, inducing double-stranded DNA breaks that can result in precise deletions, single indels, double indels, or inaccurate deletions depending on the repair mechanism. (B) Dual-plasmid CRISPR/Cas9 workflow. *M. abscessus* is first transformed with the integrative plasmid pCas9-mScarlet (KanR) encoding inducible Cas9. This Cas9-expressing strain is then transformed with pQL033-Xsg, a plasmid carrying the dual-sgRNA cassette and ZeoR selection marker. CRISPR system activation results in gene deletion and loss of function.

The sgRNA expression vector (pKMZeoR-sg) is a replicative plasmid conferring Zeocin resistance, with sgRNA transcription under tetracycline repressor control. To validate whether dual-sgRNA design could mediate gene knockout in *M. abscessus*, we targeted the *Mab_0673-Mab_0674* (*phoP/phoR*) gene cluster, designing two sgRNAs flanking the locus (Fig. S2A-B). Following electroporation into *M. abscessus* harboring pCas9-mScarlet and aTc induction, no knockout mutants were detected among 69 colonies screened (Fig. S2C). Sequencing of PCR amplicons from five clones confirmed incomplete cleavage, suggesting inefficient editing. We hypothesized that tandem arrangement of the two sgRNA cassettes might promote intermolecular or intramolecular recombination, leading to sgRNA cassette loss. Indeed, PCR and sequencing of the sgRNA region in three randomly selected clones confirmed extensive deletions in the sgRNA cassettes (Fig. S2D).

To overcome this limitation, we redesigned the sgRNA expression vector (pQL033-*Mab_0673/0674*sg), by placing the two sgRNA cassettes between the Mycobacterium replication origin (oriM) and the antibiotic resistance gene (Fig. S1, S2E). Electroporation of this modified construct into *M. abscessus* carrying pCas9-mScarlet resulted in significantly improved editing efficiency. PCR analysis 72 colonies showed that 52 colonies had successful *Mab_0673/0674* deletion, 8 lacked deletions, 1 exhibited partial deletion, and 11 produced no PCR amplicon— suggesting larger-than-expected genomic deletions (Fig. S2F).. These findings establish dual-sgRNA CRISPR/Cas9 as an effective strategy for precise gene deletion in *M. abscessus*.

### 3.2 Validation of Dual-sgRNA System Efficiency in *M. abscessus*

To further assess knockout efficiency, we targeted four additional loci (*Ms1 ncRNA, nucS, Mab_2999c*, and *Mab_2300/2301*) with designed deletions ranging from ∼200 bp to 3.5 kb (Fig. 2A,D,G,J). Deletion efficiency was consistently high but varied based on genomic context rather than fragment size.

**Figure 2.**
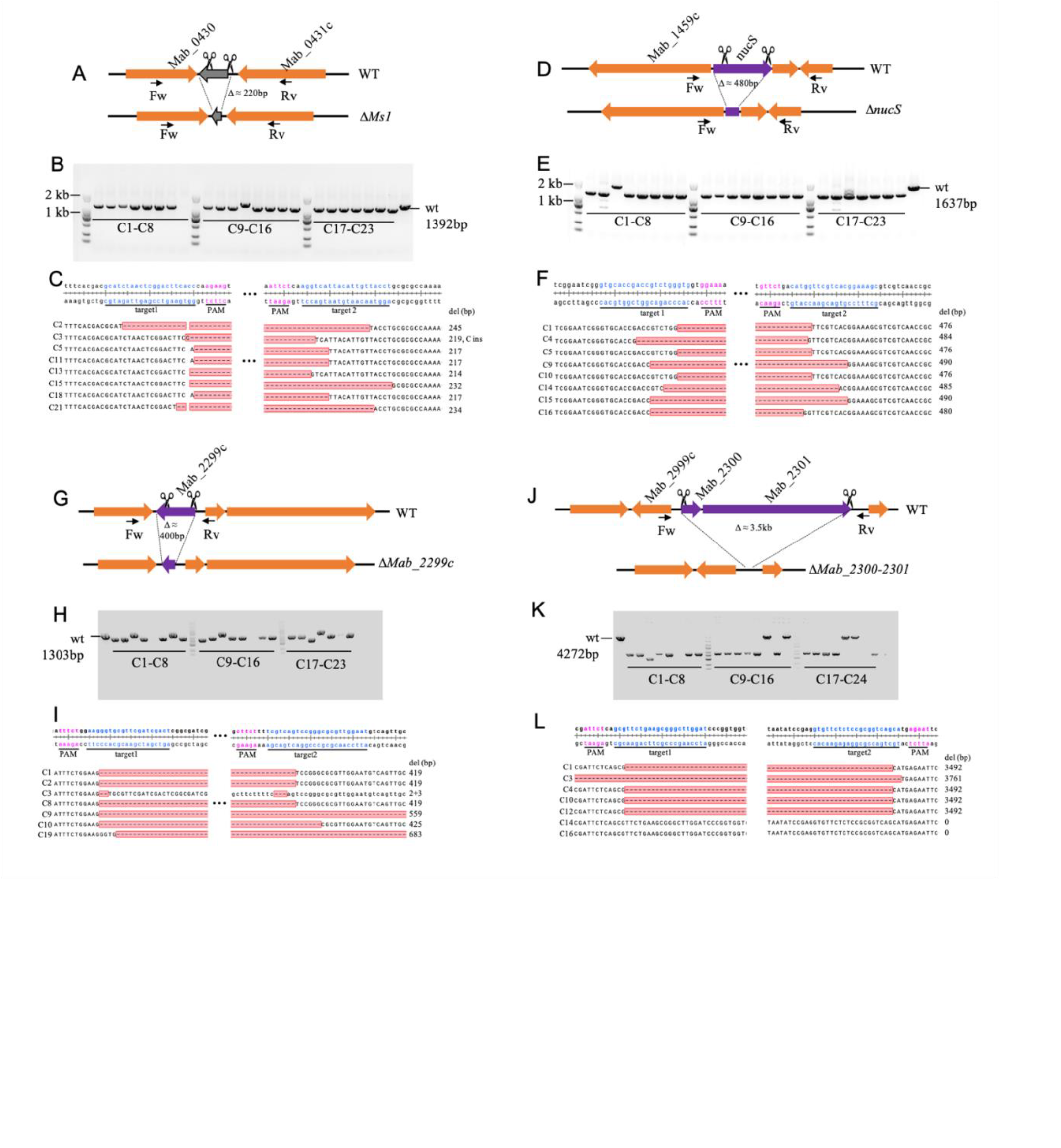
CRISPR-Cas9-mediated deletion of the *Ms1 ncRN*A, *nucS, Mab_2999c*, and *Mab_2300/2301* in *M. abscessus*. (A, D, G, J) Schematic representation of the deletions of *Ms1 ncRNA, nucS, Mab_2999c*, and *Mab_2300/2301*, respectively. (B, E, H, K) PCR verification of gene knockouts, assessed by agarose gel electrophoresis. For each knockout strain, 23 red colonies were randomly selected for colony PCR validation. Primer pairs used for verification are listed in Table S1. (C, F, I, L) Sanger sequencing confirmation of deletions in the target genomic regions, with 7–8 colonies analyzed per gene.

For *Ms1ncRNA* and *nucS* (∼200 bp and ∼480 bp deletions, respectively), correct knockout rates reached 96% (22/23 clones each) with sequencing confirming precise edits (Fig. 2B-F). In contrast, the ∼400-bp *Mab_2999c* deletion showed a lower success rate (65%, 15/23 clones), with some exhibiting additional deletions (+140 bp or +264 bp) or loss of PCR amplification (Fig. 2G-I).

Similarly, the ∼3.5 kb deletion of *Mab_2300/2301* was successful in 71% (17/24) of clones, though some exhibited larger deletions (+269 bp) or CRISPR escape events (Fig. 2J-L). These results confirm the robustness of the dual-sgRNA system while highlighting potential context-dependent variability in editing efficiency.

### 3.3 Instability of the Sth1Cas9 Expression Plasmid

During attempts to delete the methionine synthase gene *metH*, the cyclic di-AMP riboswitch (*cdAMPRibo*, the promoter region of MAB_0869c), and SRP_RNA coding sequence in *M. abscessus*, we observed high rates of pCas9-mScarlet plasmid loss following aTc induction, with white colonies appearing on selective plates (57.1%, 67.8%, and 20.7%, respectively)(Fig.3 A). PCR verification of 23 red colonies revealed no correct knockouts for *metH* and *SRP_RNA*, while only 11/23 clones showed successful *cdAMPRibo* deletion (Fig.3 B-C).

**Figure 3.**
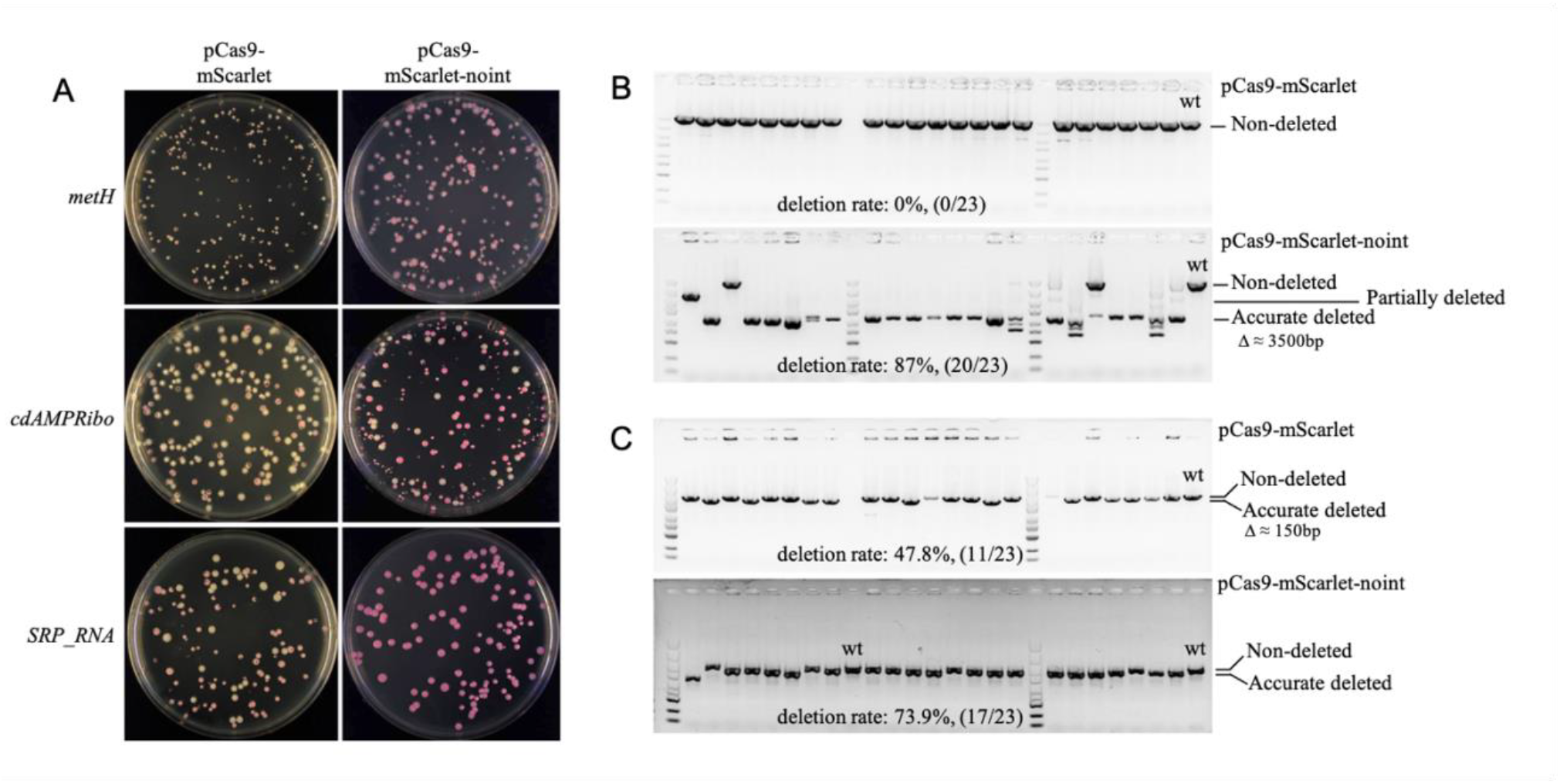
Instability of the pCas9-mScarlet plasmid in *M. abscessus*. (A) Loss of pCas9-mScarlet or pCas9-mScarlet-noint plasmids during knockout of *metH, cdAMPRibo*, and *SRP_RNA* genes. (B, C) Deletion efficiency of *metH* (B) and *cdAMPRibo* (C) validated by colony PCR and agarose gel electrophoresis. For each gene, results are shown for knockouts using the pCas9-mScarlet plasmid (top) and the pCas9-mScarlet-noint plasmid (bottom).

Although L5 integration vectors are generally stable in *M. smegmatis* (13), rare excision events can occur in the presence of L5 integrase. To prevent plasmid loss, we constructed pCas9-mScarlet-noint by removing the L5 integrase gene. Co-electroporation with pBSint (expressing transient L5 integrase) enabled integration of pCas9-mScarlet-noint without persistent integrase expression. This strategy significantly reduced plasmid loss, eliminating white colonies in *metH* and *SRP_RNA* knockouts and reducing loss to 15.5% for *cdAMPRibo* (Fig.3 A). PCR verification of randomly selected red surviving clones showed markedly improved knockout efficiency for *metH* and *cdAMPRibo*, reaching 87% (20/23) and 73.9% (17/23) respectively (Fig.3 B-C), demonstrating the importance of stabilizing Cas9 expression in some *M. abscessus* genes editing.

### 3.4 Knockout of Ultra-Long Genomic Fragments in *M. abscessus*

To evaluate the system’s capability for large-scale deletions, we targeted the *mps1-mps2* glycopeptidolipid biosynthetic cluster (18.1 kb), a non-essential region for *in vitro* growth (Fig. 4A). Following aTc induction, all surviving clones exhibited rough colony morphology, indicative of *mps1/mps2* inactivation (Fig. 4C). PCR screening of 23 rough colonies confirmed successful deletions in 43.5% (10/23), with sequencing verifying an average excision of 16.7 kb (Fig. 4B,D).

**Figure 4.**
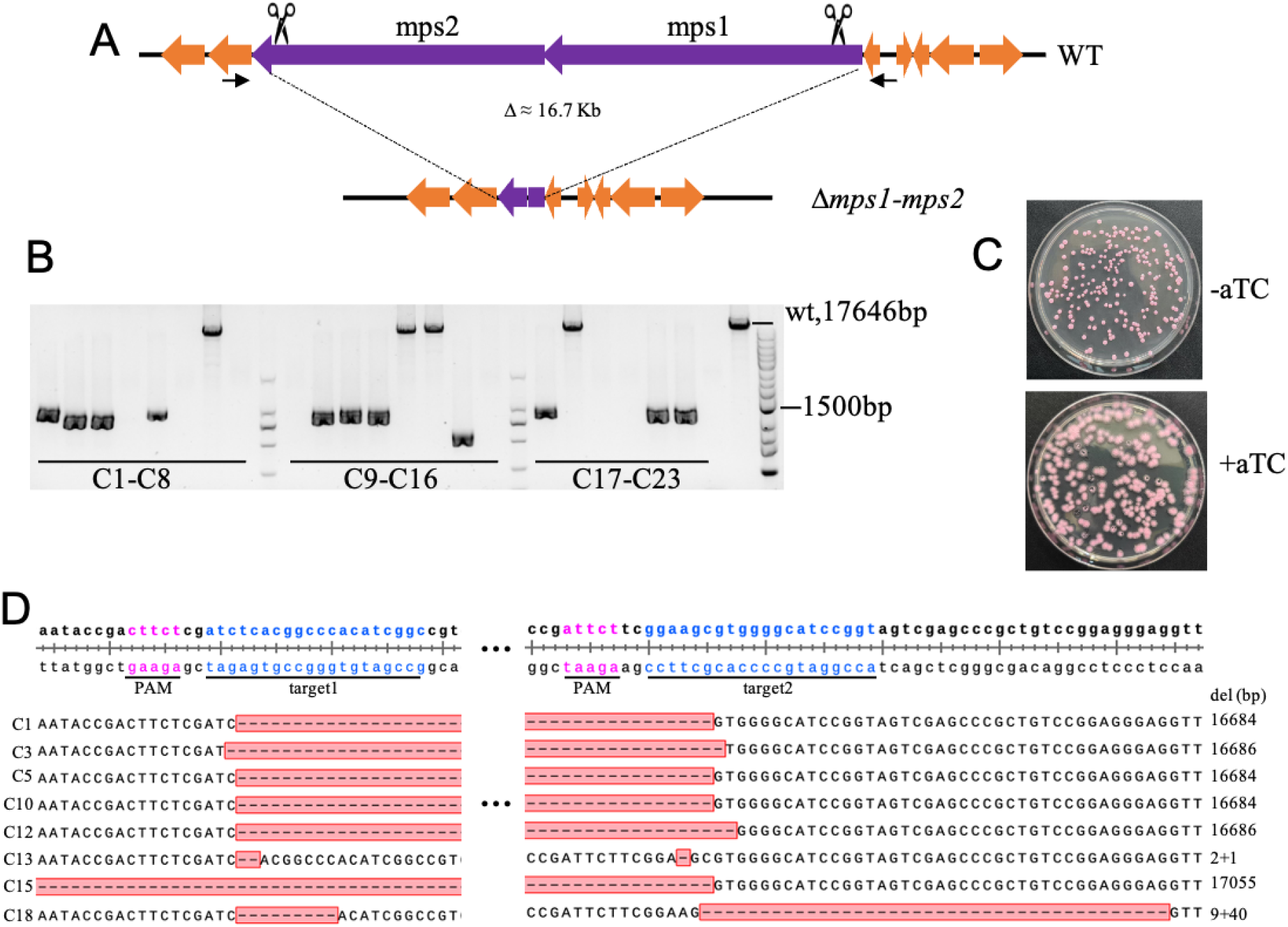
CRISPR-Cas9-mediated deletion of the 16.7-kb *mps1-mps2* fragment in *M. abscessus*. (A) Schematic representation of the *mps1-mps2* gene cluster deletion. (B) Deletion efficiency validated by colony PCR and agarose gel electrophoresis; 23 red colonies were randomly selected for analysis. (C) Colony morphology of *M. abscessus* strains carrying pCas9-mScarlet-noint and pQL033-*mps1-mps2*sg plasmids on plates with or without aTc induction. (D) Sanger sequencing confirmation of deletions in the *mps1-mps2* region from eight colonies.

One clone exhibited an additional 370 bp deletion, while eight yielded no PCR products, suggesting larger deletions affecting primer binding sites. Notably, four clones retained wild-type PCR band sizes, and sequencing revealed small deletions near sgRNA target sites (Fig. 4D). These findings establish the efficacy of dual-sgRNA CRISPR/Cas9 for ultra-large fragment deletions in *M. abscessus*.

## 4. Discussion

Current recombination-based gene editing methods exhibit low efficiency in *M. abscessus* due to its poor electroporation efficiency and high spontaneous resistance mutation rates. Recent studies have demonstrated that the Sth1Cas9 single-sgRNA system can induce targeted DNA cleavage in *M. abscessus*, leading to error-prone non-homologous end joining (NHEJ) repair that generates small indels capable of inactivating target genes through frameshift mutations (8, 10). Building upon this foundation, our study establishes a dual-sgRNA system capable of efficiently mediating large-fragment deletions in *M. abscessus*, achieving success rates exceeding 40% for deletions larger than 15 kb.

A key challenge encountered was the instability of tandem sgRNA expression cassettes, which were prone to intramolecular and intermolecular homologous recombination, leading to loss of sgRNA sequences. This issue was mitigated by repositioning the sgRNA cassettes between the plasmid replication origin (*oriM*) and the zeocin resistance gene (*ZeoR*). Furthermore, we developed intermediate plasmids (pQL033-SapItwin-sgSC and pUC19-CS-ZeoR) to facilitate one-step assembly of dual-sgRNA expression vectors, streamlining cloning procedures and enabling efficient CRISPR knockout plasmid library construction (Fig. S1).

Interestingly, we observed no linear inverse correlation between deletion efficiency and fragment size. For instance, deletion of the 3.5-kb *Mab_2300/2301* locus achieved 71% efficiency, whereas deletion of the 150-bp *cdAMPRibo* locus resulted in only 47.8% efficiency. As the protospacers had comparable PAM sequence strength and GC content, this variability is likely due to locus-specific differences in double-strand break (DSB) repair efficiency or spatial constraints between break ends. Future studies targeting additional loci and analyzing larger mutant pools will be required to elucidate the underlying mechanisms influencing deletion efficiency.

To mitigate the cytotoxic effects of simultaneous Cas9 and sgRNA expression—a phenomenon reported previously (10)—we separated Cas9 and sgRNA expression into distinct plasmids for sequential transformation. Additionally, an mScarlet reporter, constitutively expressed from the P_left_ promoter, was integrated into the Sth1Cas9 genomic expression plasmid, allowing real-time monitoring of plasmid retention. Notably, certain knockouts, including *metH* and *cdAMPRibo*, resulted in approximately 60% of colonies turning white after aTc induction, suggesting frequent loss of the pCas9-mScarlet plasmid. Red colonies, indicative of plasmid retention, exhibited minimal knockout success, implying a strong counterselection effect at these loci. Removal of the L5 integrase from the Cas9 expressing plasmid significantly reduced the proportion of white colonies, confirming that integrase-mediated excision was the primary loss mechanism. However, residual white colonies in cdAMPRibo knockouts, even with integrase-deficient plasmids, suggest the presence of additional plasmid loss or inactivation pathways. These findings underscore the necessity of mitigating Cas9 plasmid loss when employing Sth1Cas9 for *M. abscessus* genetic manipulation, particularly for constructing CRISPR-based mutant libraries. Strategies such as using integrase-deficient integration plasmids are crucial for maintaining plasmid stability, albeit potentially at the cost of reduced electroporation efficiency when supplemental integrase plasmids are required.

Importantly, our study found that *M. abscessus* exhibits low spontaneous resistance to zeocin, suggesting the utility of ZeoR-based selection systems for future genome-editing applications.

In conclusion, we have established a robust, dual-sgRNA CRISPR/Cas9 system for efficient large-fragment deletions in *M. abscessus*. This streamlined, helper-factor-free approach provides a powerful genetic tool for studying bacterial gene function, pathogenicity, and antibiotic resistance in this clinically significant and genetically recalcitrant pathogen.

