## Supplementary material for "A CRISPR/Cas9-based system using dual-sgRNAs for efficient gene deletion in *Mycobacterium abscessus*": Fig.S1, Fig.S2, Table S1

### 1 Supplementary Data

Supplementary Material should be uploaded separately on submission. Please include any supplementary data, figures and/or tables.

Supplementary material is not typeset so please ensure that all information is clearly presented, the appropriate caption is included in the file and not in the manuscript, and that the style conforms to the rest of the article.

### 2 Supplementary Figures and Tables

For more information on Supplementary Material and for details on the different file types accepted, please see here.

#### 2.1 Supplementary Figures

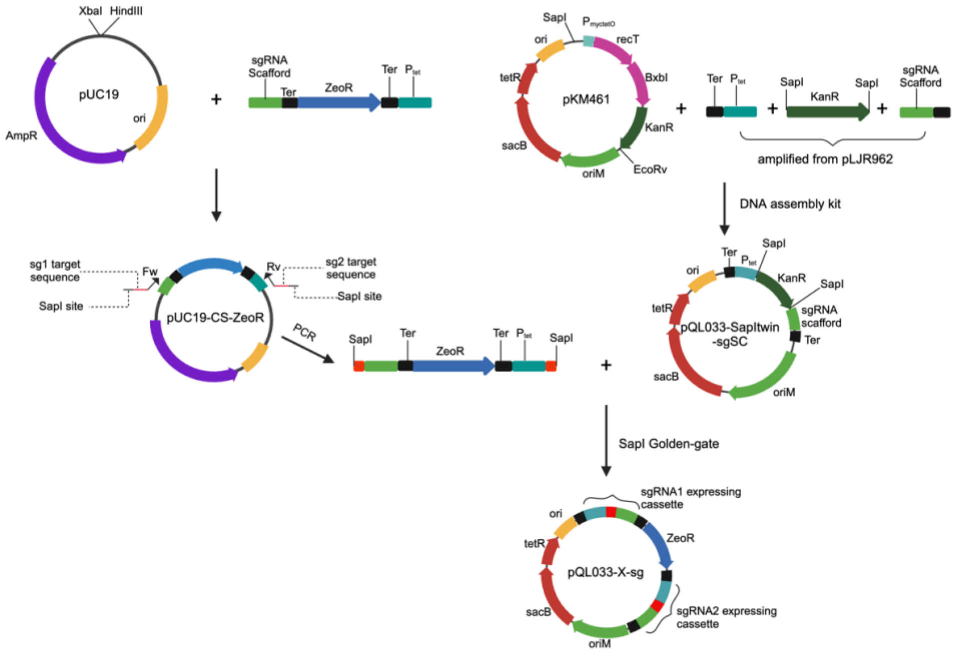


**Fig.S1.** Schematic of the dual-sgRNA-expressing pQL033-Xsg plasmid construction.

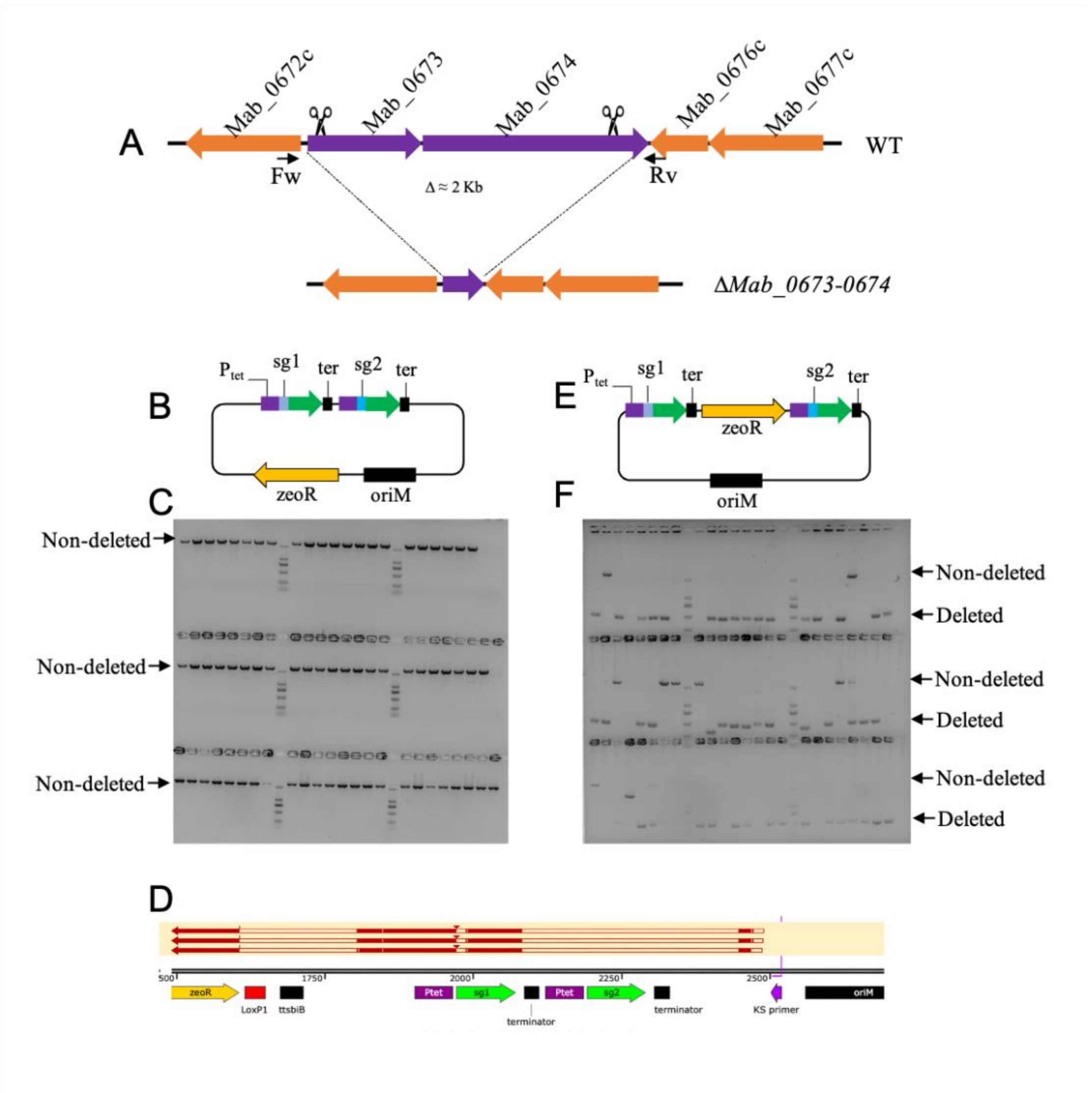


**Fig. S2. Comparative analysis of editing efficiencies across dual-sgRNA expression plasmids**

(A) Schematic representation of *Mab_0673/0674* gene cluster knockout. (B) Plasmid architecture schematic of pKMZeoR-*Mab_0673/074*-sg construct. (C) Colony PCR and agarose gel electrophoresis validation of *Mab_0673/074* knockout in *M. abscessus* strains harboring pCas9-mScarlet and pKMZeoR-*Mab_0673/0674*-sg plasmids following aTc induction.. (D) Loss of sgRNA expression cassette in pKMZeoR-*Mab_0673/0674*-sg plasmid, assessed by PCR amplification and Sanger sequencing of the sgRNA cassette region in three randomly selected clones from panel C. (E) Plasmid architecture of pQL033-*Mab_0673/074*-sg construct. (F) Colony PCR and agarose gel electrophoresis validation of *Mab_0673/0674* knockout in *M. abscessus* strains carrying pCas9-mScarlet and pQL033-*Mab_0673/0674*-sg plasmids after aTc induction.

### 2.2 Supplementary Tables

**Table S1. Oligonucleotides used in this study**. Restriction sites are underlined (SapI), OVERhang regions are italicized, CRISPR-Cas9 sgRNA targeting regions are Bolded.

| # | Primer name | Sequence(5' → 3') |
|---|---|---|
| 1 | MabHeesg1_FW_SapIGGA | GGCTACGCTCTTCG*GGA***AGGTAACAATGTAATGACCT**GTTTTGTACTCGAAAGAAGCTACAAAGA |
| 2 | MabHeesg2_Rv_SapIGTT | GGCTACGCTCTTCG*AAC***GGTGAAGTCCGAGTTAGATGC**TCCCAGATTATATCTATCACTGATAGGGAT |
| 3 | MabnucSsg1_FW_SapIGGA | GGCTACGCTCTTCG*GGA***GTGCACCGACCGTCTGGGTG**GTTTTGTACTCGAAAGAAGCTACAAAGA |
| 4 | MabnucSsg2_Rv_SapIGTT | GGCTACGCTCTTCG*AAC***CATGGTTCGTCACGGAAAGC**TCCCAGATTATATCTATCACTGATAGGGAT |
| 5 | MabNcRNA1sg1_FW_SapIGGA | GGCTACGCTCTTCG*GGA***ACCTCGCAGGTGCGTCACAGA**GTTTTGTACTCGAAAGAAGCTACAAAGA |
| 6 | MabNcRNA1g2_Rv_SapIGTT | GGCTACGCTCTTCG*AAC***CGCGGTCCACACCCGAGGGAC**TCCCAGATTATATCTATCACTGATAGGGAT |
| 7 | MabNcRNA2sg1_FW_SapIGGA | GGCTACGCTCTTCG*GGA***ACCCCCACCACAGTTAACGCTGGGG**GTTTTGTACTCGAAAGAAGCTACAAAGA |
| 8 | MabNcRNA2g2_Rv_SapIGTT | GGCTACGCTCTTCG*AAC***AGATCGCGCCTCCGGATGCC**TCCCAGATTATATCTATCACTGATAGGGAT |
| 9 | MabmetHsg1_FW_SapIGGA: | GGCTACGCTCTTCG*GGA***AGTTAAGGACGCCTTCCGCT**GTTTTGTACTCGAAAGAAGCTACAAAGA |
| 10 | MabmetHsg2_Rv_SapIGTT | GGCTACGCTCTTCG*AAC***GTGTGCTCGGGGCAGGCCGGGT**TCCCAGATTATATCTATCACTGATAGGGAT |
| 11 | Mab_2299csg1_FW_SapIGGA | GGCTACGCTCTTCG*GGA***AGTCGATCGAACGCACCCTT**GTTTTGTACTCGAAAGAAGCTACAAAGA |
| 12 | Mab_2299csg2_Rv_SapIGTT | GGCTACGCTCTTCG*AAC***ATTCCAACGCGCCCGGACTGACGA**TCCCAGATTATATCTATCACTGATAGGGAT |

| | | |
|---|---|---|
| 13 | Mab_2300_2301sg1_FW_SapIGGA | GGCTACGCTCTTCG*GGA***ATCCAAGCCCGCTTCAGAACGC**GTTTTTGTACTCGAAAGAAGCTACAAAGA |
| 14 | Mab_2300_2301sg2_Rv_SapIGTT | GGCTACGCTCTTCG*AAC***TGCTGACCGCGGAGAGAACAC**TCCCAGATTATATCTATCACTGATAGGGAT |
| 15 | Mab_0673sg1_FW_SapIGGA_1: | **GCGGGGGTGGTGCCGTTGGC**GTTTTTGTACTCGAAAGAAGCTACAAAGA |
| 16 | Mab_0673sg1_FW_SapIGGA_2 | GGGCTACGCTCTTCG*GGA***GCGGGGGTGGTGCCGTTGGC**GTT |
| 17 | Mab_0674sg2_Rv_SapIGTT_1 | **CGCGCGGATTCCTCGCGCAC**TCCCAGATTATATCTATCACTGATAGGGAT |
| 18 | Mab_0674sg2_Rv_SapIGTT_2 | GGCTACGCTCTTCG*AAC***CGCGCGGATTCCTCGCGCAC**T |
| 19 | MabMps1sg1_FW_SapIGGA_1 | **ACCGGATGCCCCACGCTTCC**GTTTTTGTACTCGAAAGAAGCTACAAAGA |
| 20 | MabMps1sg1_FW_SapIGGA_2 | GGCTACGCTCTTCG*GGA***ACCGGATGCCCCACGCTTCC**GTT |
| 21 | MabMps2sg2_Rv_SapIGTT_1 | **ATCTCACGGCCCACATCGGC**TCCCAGATTATATCTATCACTGATAGGGAT |
| 22 | MabMps2sg2_Rv_SapIGTT_2 | GGCTACGCTCTTCG*AAC***ATCTCACGGCCCACATCGGC**T |
| 23 | MABHeeKOcon_FW | ATGGCAGATGAGGACACC |
| 24 | MABHeeKOcon_Rv | ATGTGGAACAAGTGAAGGC |
| 25 | MABnucSKOcon_FW | GGACTATGACGAAGTTGAC |
| 26 | MABnucSKOcon_Rv | GAGGCCATGCTGTCCTTT |

| | | |
|---|---|---|
| 27 | MABNcRNA1KOcon_FW | ATCGACGAGATGGCTGTG |
| 28 | MABNcRNA1KOcon_Rv | AGCATCACGGTCAGGTAT |
| 29 | MABmetHKOcon_FW | AGCAAGCGGTTGCAGAAC |
| 30 | MABmetHKOcon_Rv | CCATCGCACCAGATGAGA |
| 31 | MAB2299cKOcon_FW | TTAGCGGGCATCGGGTTG |
| 32 | MAB2299cKOcon_Rv | CGAACCTGGCTCAACTACCG |
| 33 | MAB2300_2301KOcon_FW | GCGTTGGAATGTCAGTTGCG |
| 34 | MAB2300_2301KOcon_Rv | GTTCACCGCCCTAAGCAC |
| 35 | MAB0673_74KOcon_FW | CCAATCCGGTGTCGGCAA |
| 36 | MAB0673_74KOcon_Rv | CGCACGATCTTTGGTTCGGA |
| 37 | MABmps1_2KOcon_FW | GATTCATCACGTGGTAGGTCTC |
| 38 | MABmps1_2KOcon_Rv | GTTCGGATCGAAGGTCAAGTAG |
